# Viscosity-mediated signal amplification enables robust left–right symmetry breaking in the mouse node

**DOI:** 10.64898/2026.01.13.699291

**Authors:** Chiu Fan Lee, Julien Vermot

## Abstract

Left–right (LR) symmetry breaking in vertebrates depends on a directional fluid flow generated in the embryonic LR organizer, yet how this flow is sensed remains unresolved. Mechanosensing and chemosensing models each capture part of the process but face key limitations when considered independently. Here, we introduce a minimal theoretical model of viscosity-mediated signal amplification that unifies these perspectives. In this framework, macromolecules secreted by organizer cells locally increase the near-surface viscosity, creating another viscous fluid layer that is entrained by the leftward nodal flow. This layer can significantly amplifies the drag and torque exerted on immotile perinodal cilia, enabling robust discrimination of flow direction even when flow magnitudes on the left and right are nearly identical. The model naturally incorporates macromolecule secretion, clarifies the complementary roles of motile and immotile cilia, and resolves the major shortcomings of previous proposals. Together, these results provide a simple and physically grounded mechanism for LR determination in the mouse node.

## I. INTRODUCTION

Chirality—an asymmetry between an object and its mirror image—has long captivated researchers across physics and biology [1, 2]. In living systems, it emerges at every scale: from chiral cortical flows in single cells [3] to tissue-level processes such as left–right (LR) axis determination during embryonic development [4–8]. Similar principles operate in plants as well, underscoring the ubiquity and biological significance of chirality [9].

Biological symmetry breaking frequently arises from the intrinsic handedness of molecular motors and cytoskeletal assemblies [10–12]. Among the most striking examples is LR axis determination in vertebrate embryos, which ensures the asymmetric positioning of internal organs. In these systems, LR specification is initiated by a symmetry-biasing event produced by the chiral beating of motile cilia. These cilia generate a unidirectional fluid flow at the left–right organizer (LRO)—the embryonic structure where LR determination takes place [13, 14]. Although the downstream signaling cascade (Nodal–Lefty–Pitx2) is conserved across vertebrates [15], the upstream mechanism by which flow is sensed remains incompletely understood [10, 16].

In the mouse, LR symmetry breaking occurs at the node, a transient LRO composed of centrally located pit cells with motile cilia and surrounding crown (perinodal) cells bearing immotile, “sensory” cilia. Tilted, clockwise-rotating motile cilia (∼200 nm in diameter and ∼4 *µ*m long) generate a robust leftward flow across the node [17]. While this flow is known to be essential for LR specification [18, 19], how it is detected by the perinodal cells remains controversial.

Two main classes of hypotheses have been proposed for flow sensing. (I) The mechanosensing model posits that sensory cilia on perinodal cells detect flow-induced mechanical stresses, potentially via mechanosensitive calcium channels such as Pkd2 [20, 21]. (II) The chemosensing model suggests that leftward flow transports signaling macromolecules, such as nodal vesicular parcels, preferentially to the left margin, where they are absorbed and initiate downstream signaling [18].

Both mechanisms have empirical support, yet they are often treated as mutually exclusive. Here, we first highlight the key limitations of each model when considered in isolation and then present a minimal biophysical framework that integrates elements of both, demonstrating how their synergy can robustly account for LR determination in the mouse node.

## II. LIMITATIONS OF THE TWO EXISTING MODELS

We now highlight the key limitations of each proposal when considered in isolation.

### Issues with proposal I (mechanosensing)

**I.1:** Analytical, computational, and experimental studies have shown that fluid-flow magnitudes above immotile cilia are nearly identical on the left and right sides of the node (Fig. 1a) [22– Consequently, the mechanical stresses transmitted to immotile cilia on either side should also be comparable, making it unclear how such stresses alone could provide a robust mechanism for LR asymmetry.

**I.2:** Fluid-dynamic analyses indicate that the flow generated by a single rotating cilium decays with the cube of the distance [27]. Thus, the flow experienced by any given peripheral crown cell is dominated by contributions from only the nearest rows of motile cilia (Fig. 1a). If flow magnitude were the sole relevant factor, it would be puzzling why the mouse node contains a large population of motile cilia concentrated at its center.

**FIG. 1.**
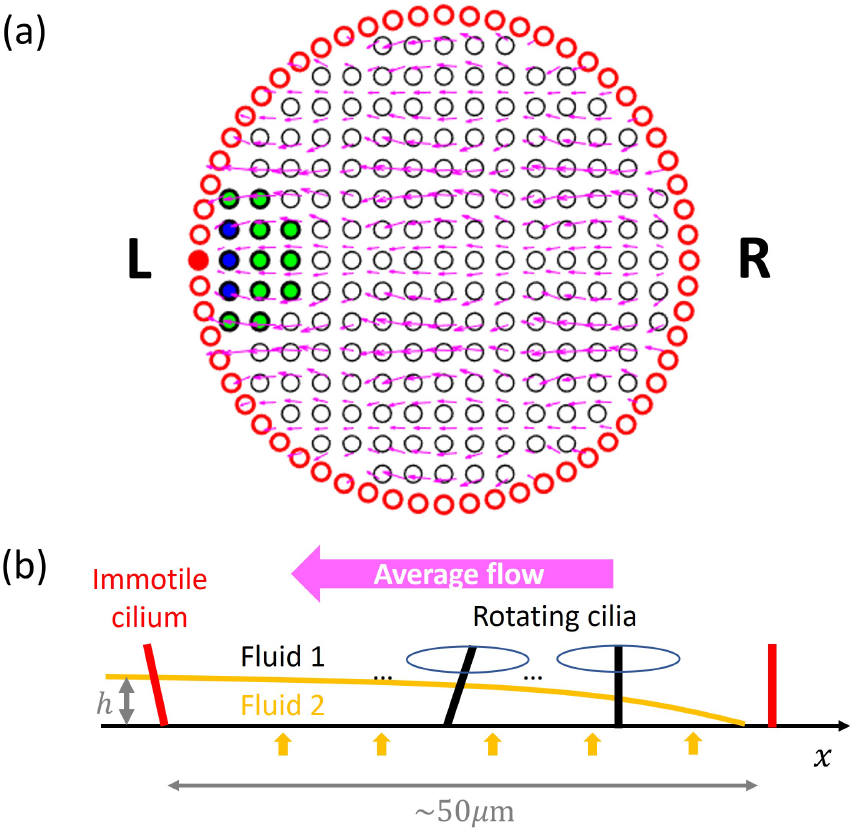
Schematics of the mouse node, the organ that establishes organismal left–right asymmetry. (a) *Top view*. The mouse node (∼50 *µ*m in diameter) is modeled as 177 cells (∼ 3 *µ*m each, black circles) bearing motile cilia that are tilted and rotate clockwise, collectively generating a leftward fluid flow (magenta arrows). These motile cilia are surrounded by crown cells bearing immotile “sensory” cilia (red circles), which detect the resulting flow. Because the flow produced by each motile cilium decays rapidly with distance, the local flow at any immotile cilium is dominated by nearby sources. For example, for the leftmost red cell (highlighted), 82% of the flow arises from the three nearest blue cells, and 98% from the blue and green cells combined. (b) *Side view*. Pit cells bearing motile cilia (black) occupy the center of the node, while immotile cilia (red) reside on surrounding crown cells. Recent experiments show that nodal cells secrete macromolecules (orange arrows). In our model, these secretions give rise to a near-surface high-viscosity layer (Fluid 2, thickness *h*) adjacent to the tissue on the left side. This viscous layer is entrained by the leftward flow generated by motile cilia and increases the drag experienced by left immotile cilia relative to the right.

### Issues with proposal II (chemosensing)

**II.1:** Node cells are known to secrete macromolecules, including polymers and vesicles [19, 28, 29]; however, none have been identified as bona fide signaling molecules for LR determination.

**II.2:** This proposal does not assign a role to immotile cilia. Yet experiments show that LR determination can be triggered by mechanically pulling on immotile cilia [30, 31], strongly suggesting that mechanical stress on these cilia is integral to the sensing mechanism.

Taken together, these limitations indicate that neither proposal alone can fully account for LR symmetry breaking. We now show how a reliable LR symmetry–determining mechanism emerges by integrating key elements of both models.

## III. A TWO-FLUID MODEL

Our model begins from three well-established experimental observations: 1) nodal cells secrete macromolecules into the fluid environment; 2) an approximately constant leftward flow exists above the cilia; and 3) mechanical pulling of immotile cilia on peripheral crown cells is sufficient to trigger the downstream signaling cascade specifying the “left” direction. Building on this foundation, we introduce two additional assumptions:

**Assumption 1:** *High-viscosity fluid layer*. Secreted macromolecules increase the viscosity of the fluid immediately adjacent to the tissue surface. In our minimal model, we represent this region as a distinct fluid phase (Fluid 2, Fig. 1b) with higher viscosity than the bulk fluid (Fluid 1) that fills the rest of the node. Although these fluids may not be perfectly unmixed *in vivo*, this simplification captures the essential physics, and the resulting qualitative behavior should remain robust.

**Assumption 2:** *Coupled two-phase flow*. Fluid 2 is entrained leftward by the overlying flow of Fluid 1 generated by motile cilia.

When combined, these ingredients give rise to a key emergent behavior: *macromolecules secreted by nodal cells locally increase near-surface viscosity, thereby capable of amplifying the mechanical stresses imposed by left-ward flow on immotile cilia*. As shown below, this “viscous amplification” mechanism reconciles existing observations and provides a simple, robust basis for distinguishing left from right during embryonic development.

To elucidate the qualitative behaviour emerging from our model, we analyse a simplified setting. (i) We reduce the system to a one-dimensional description along the left–right (LR) axis; and (ii) we quantify the flow-induced drag acting on immotile cilia located on perinodal cells at the left and right peripheries.

For the fluid dynamics, we impose boundary conditions consistent with prior computational and experimental studies: a no-slip condition at the tissue surface and a uniform leftward flow at a short distance above the cilia. The latter is implemented as a no-shear condition at height *H* = 5*µ*m above the tissue surface. For a homogeneous single-phase fluid, these conditions yield the standard half-Poiseuille velocity profile below height *H* (top panel of Fig. 2b),

**FIG. 2.**
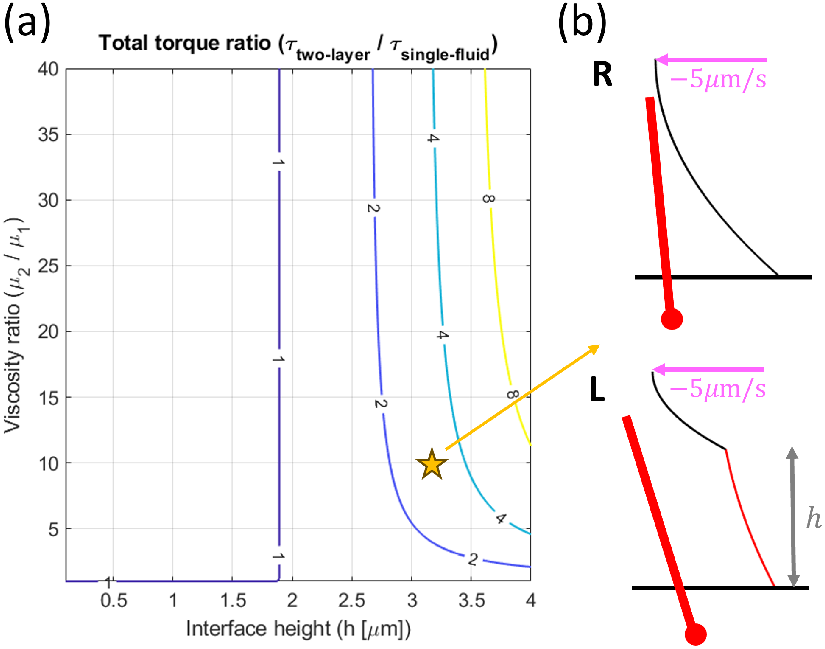
Contour plot of the ratio of total torque experienced by left versus right immotile cilia. (a) The torque ratio is plotted as a function of the viscosity ratio between the two fluid layers (vertical axis) and the thickness *h* of Fluid 2 at the left periphery of the node (horizontal axis; see Fig. 1b). In this idealized one-dimensional representation, the flow speed 5*µ*m above the tissue surface is fixed at −5*µ*m/s [23] (negative indicates leftward flow), with a no-shear boundary condition. On the right periphery, the flow reduces to a standard half–Poiseuille profile (top of (b)). On the left, a two-phase flow forms with viscosities *µ*_1_ (top layer) and *µ*_2_ (bottom layer) (bottom of (b)). Although the higher viscosity of Fluid 2 slows its flow (red profile), drag is proportional to the product of viscosity and velocity, so the net torque on left immotile cilia can exceed that on the right. For instance, when *h* = 3.2, *µ*m and *µ*_2_*/µ*_1_ = 10 (yellow star), the torque is amplified by around threefold due to the additional viscous layer, with the corresponding flow profiles shown in (b).

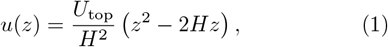

where *U*_top_ denotes the flow speed at height *H*.

When a second fluid layer is present near the tissue surface (Fluid 2, viscosity *µ*_2_), it is entrained by the left-ward flow of the overlying Fluid 1 (viscosity *µ*_1_). Assuming that the fluid–fluid interface remains approximately parallel to the tissue surface, and imposing a no-shear condition at this interface, we obtain an analytical expression for the velocity profile experienced by immotile cilia on the left side of the node (bottom panel of Fig. 2b). Although this expression is fully analytical, it is lengthy and not particularly illuminating; we therefore relegate it to the Appendix.

To quantify the mechanical signal generated by the flow, we use the torque exerted on immotile cilia as a proxy for LR-determining cues. This choice is motivated by experimental evidence that immotile cilia behave as hinged rods anchored at a short distance (∼ 1*µ*m) below the cell surface [32]. While the scenario considered here is highly idealized, we expect our central qualitative result—the existence of a viscosity-induced amplification mechanism—to be robust and largely independent of the specific modelling assumptions adopted.

## IV. RESULTS

Crucially, the model predicts a qualitative asymmetry between left and right. Immotile cilia on the right experience a uniform Poiseuille flow of Fluid 1, whereas those on the left are exposed to a two-tiered flow: a slow but highly viscous layer of Fluid 2 adjacent to the tissue surface, overlaid by the faster-moving bulk flow of Fluid 1. Although the near-surface viscous layer moves more slowly, drag depends on both velocity and viscosity. As a result, the viscosity-induced enhancement of drag and torque acting on left immotile cilia can outweigh the reduction in flow speed, leading to an overall increase in mechanical stimulation.

Figure 2 shows the ratio of the total torque acting on left versus right immotile cilia as a function of the viscosity contrast between Fluids 1 and 2 and the thickness of Fluid 2 at the left periphery. Because the flow profiles are analytically tractable, one can show explicitly that when the height of Fluid 2 exceeds a critical threshold, increasing its viscosity enhances the torque on left immotile cilia relative to the right (see Appendix). This generates a robust and reliable LR-asymmetric mechanical signal, whose amplification grows nonlinearly with either increasing viscosity contrast or increasing Fluid 2 thickness.

Beyond providing a robust physical mechanism for LR determination, the model naturally resolves the limitations of existing proposals. First, it is consistent with analytical, computational, and experimental evidence showing that flow magnitudes above immotile cilia are nearly identical on the left and right, thereby resolving issue I.1. Second, it clarifies the role of the large population of centrally located motile cilia: their collective action is required both to sustain the leftward flow and to secrete macromolecules that generate the viscous near-surface layer, resolving issue I.2. Third, the model obviates the need to invoke unidentified signaling molecules; in our framework, secreted macromolecules merely increase the effective near-surface viscosity, and their molecular identity is irrelevant for LR determination, resolving issue II.1. Finally, the model highlights the central role of immotile cilia as mechanosensors that detect the amplified stresses—particularly torque—produced by the viscous layer, thereby providing the decisive cue for symmetry breaking, resolving issue II.2.

## V. SUMMARY & OUTLOOK

Our two-fluid model provides a minimal, physically grounded framework for left–right (LR) determination in the mouse node, unifying mechanosensing and chemosensing into a coherent mechanism for symmetry breaking. In this picture, immotile cilia sense torque generated by a near-surface viscous layer. Secreted macromolecules do not act as signaling ligands; rather, they locally increase viscosity, thereby amplifying the mechanical stresses imposed by leftward flow. This “viscous amplification” enables reliable discrimination between left and right even when flow magnitudes are nearly identical.

The model also clarifies the role of the centrally located motile cilia, which both sustain the leftward flow and maintain the viscous layer through secretion. Because the mechanism relies only on directional bias—not absolute flow speed—LR symmetry breaking can in principle occur with remarkably few motile cilia [34, 35]. Robustness emerges as another key feature: the mechanism is largely insensitive to cilia length, the precise molecular identity of secreted components, and variations in flow strength, making it resilient to biological variability and noise.

Taken together, the viscous-amplifier model reconciles the strengths of the two leading proposals and offers a simple, experimentally consistent mechanism for LR symmetry breaking in the mouse node. A promising next step will be to probe experimentally the amount, composition, and spatial distribution of secreted macromolecules, in order to quantify the viscosity and thickness of the high-viscosity layer more precisely.

## Appendix

### TWO-PHASE (TWO-LAYER) STOKES FLOW: DERIVATION OF VELOCITY PROFILES AND TOTAL TORQUE

#### PHYSICAL SETUP

We consider steady, planar, laminar flow of two immiscible Newtonian layers of total height *H*, with the interface located at *z* = *h* (0 < *h* < *H*). The bottom layer occupies 0 ≤ *z* ≤ *h* and has viscosity *µ*_2_, while the top layer occupies *h* ≤ *z* ≤ *H* and has viscosity *µ*_1_. We nondimensionalise by the top-layer viscosity, setting *µ*_1_ ≡ 1 and defining the viscosity ratio *µ* = *µ*_2_*/µ*_1_ = *µ*_2_. The bottom wall at *z* = 0 is stationary (no slip), and the top wall at *z* = *H* moves at speed *U*.

#### Governing equations and general solution

In each layer, the steady Stokes equation reduces to

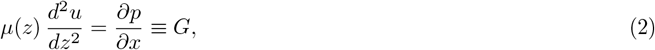

where *µ*(*z*) = *µ* for 0 ≤ *z* ≤ *h* and *µ*(*z*) = 1 for *h* ≤ *z* ≤ *H*, and *G* is constant (uniform pressure gradient). Therefore the velocitz profiles are quadratic in *z*:

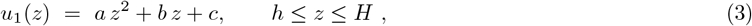

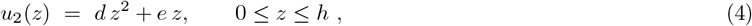

where the no-slip boundary condition at *z* = 0 has been imposed. Further, the second derivatives are

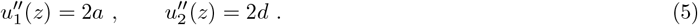

From 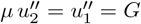 it follows that

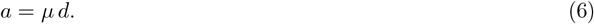

#### Boundary and interface conditions

We impose the following boundary and interface conditions:

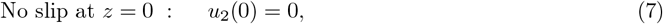

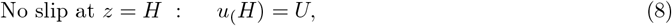

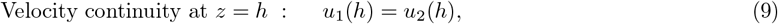

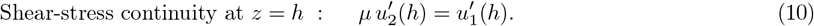

Since

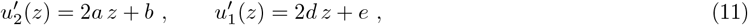

shear-stress continuity with *a* = *µd* implies

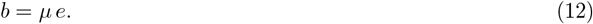

Velocity continuity determines *c* as

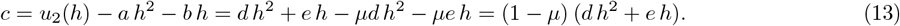

Enforcing *u*_1_(*H*) = *U* then yields a linear equation for *d* and *e*:

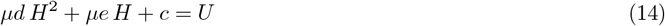

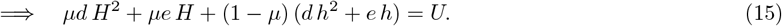

Solving for *d* and *e* gives the closed-form coefficients

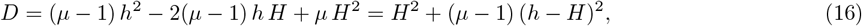

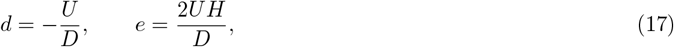

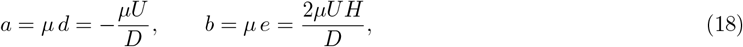

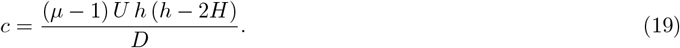

#### Two-phase velocity profiles

The distinct two-layer velocity profiles are thus

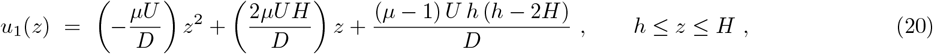

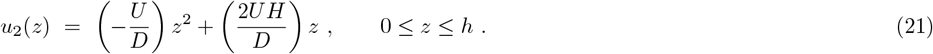

#### Total torque exerted on an immotile slender rod

We assume that the immotile cilium is almost perpendicular to the flow direction, the local hydrodynamic drag per unit length is thus dominated by transverse drag:

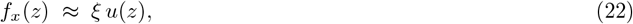

where *ξ* is the transverse drag coefficient per unit length.

For a slender rod of length *L* and radius *r* in a fluid of viscosity *µ*,

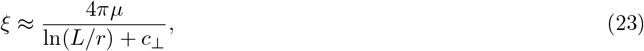

with *c*_⊥_ = 𝒪 (1) depending on the slender-body approximation and wall corrections.

In a two-layer flow, the total torque about the hinge point (at a distance ℓ below the cell surface about the *y* axis) due to the distributed force is

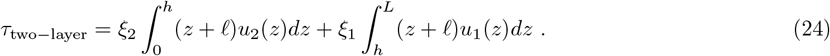

Since we are interested in the amplification effect of a two-layer flow, we will consider the ratio of the total torque in a two-fluid flow vs. that of the a single flow. Using our expression for flow profiles, the ratio *τ*_two− layer_*/τ*_single− fluid_ is

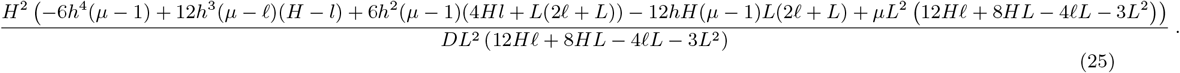

The above expression is used to compute the contour plot in Fig. 2a), with the following parameters: *H* = 5*µ*m, *L* = 4*µ*m, and ℓ = 1*µ*m. The values for *H* and *L* are well documented from existing experiments. The value for ℓ—the distance between the hinge of the cilium and the cell surface—is an estimate. Since the hinge is likely bounded by between the basal body and the nuclear membrane, we take ℓ to be 1 *µ*m [32]. However, the quantification shown in Fig. 2a) is relatively insensitive to what the actual value is, as shown in Fig. 3.

**FIG. 3.**
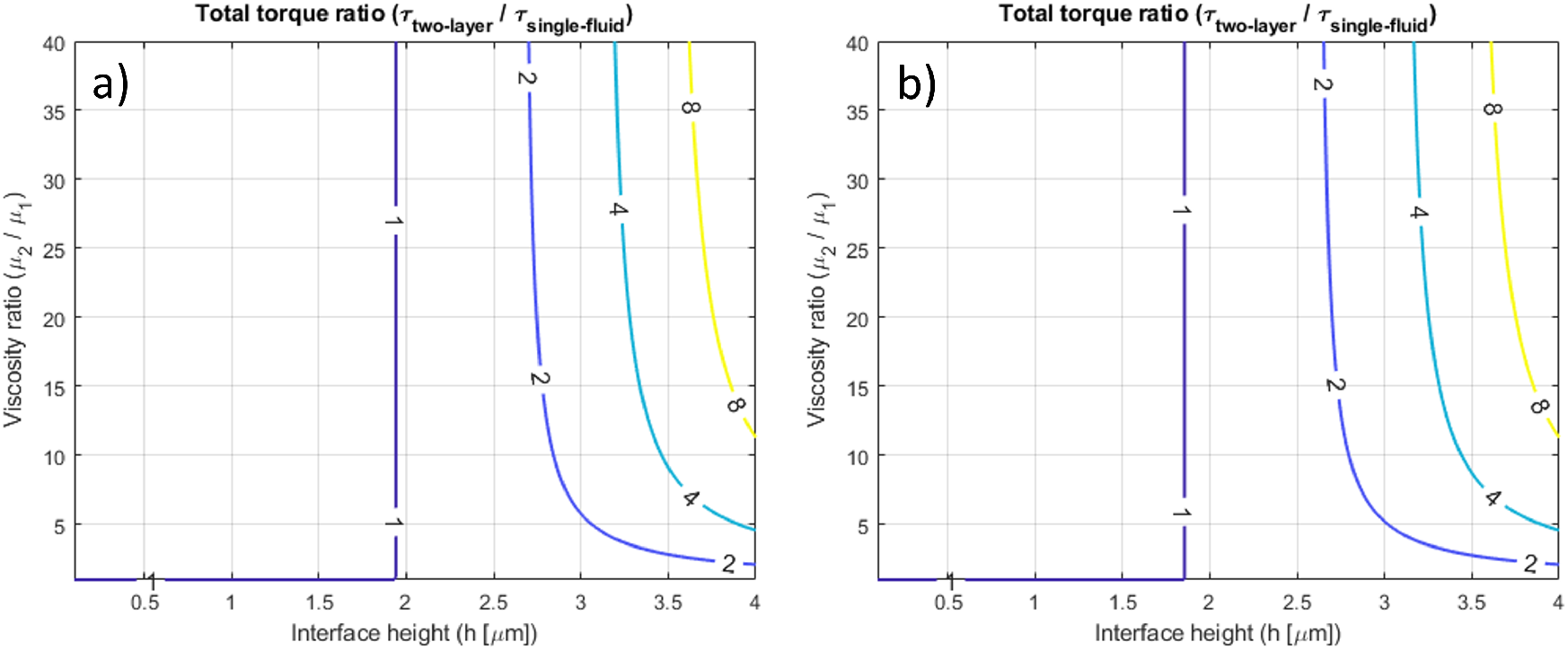
Contour plot of the ratio of total torque experienced by left versus right immotile cilia as in Fig. 2a), but for different values of ℓ—the distance of the cilium hinge from the cell surface: a) ℓ = 0.5*µ*m, b) ℓ = 1.5*µ*m.

